# Kinetochore capture by spindle microtubules: why fission yeast may prefer pivoting to search-and-capture

**DOI:** 10.1101/673723

**Authors:** Indrani Nayak, Dibyendu Das, Amitabha Nandi

## Abstract

The mechanism by which microtubules find kinetochores during spindle formation is a key question in cell biology. Previous experimental studies have shown that although search-and-capture of kinetochores by dynamic microtubules is a dominant mechanism in many organisms, several other capture mechanisms are also possible. One such mechanism reported in *Schizosaccharomyces pombe* shows that microtubules can exhibit a prolonged *pause* between growth and shrinkage. During the pause, the microtubules pivoted at the spindle pole body search for the kinetochores by performing an angular diffusion. Is the latter mechanism purely accidental, or could there be any physical advantage underlying its selection? To compare the efficiency of these two mechanisms, we numerically study distinct models and compute the timescales of kinetochore capture as a function of microtubule number *N*. We find that the capture timescales have non-trivial dependences on microtubule number, and one mechanism may be preferred over the other depending on this number. While for small *N* (as in *fission yeast*), the *typical capture times* due to rotational diffusion are lesser than those for search-and-capture, the situation is reversed beyond a certain *N*. The capture times for rotational diffusion tend to saturate due to geometrical constraints, while those for search-and-capture reduce monotonically with increasing *N* making it physically more efficient. The results provide a rationale for the common occurrence of classic search-and-capture process in many eukaryotes which have few hundreds of dynamic microtubules, as well as justify exceptions in cells with fewer microtubules.

---

An essential function of a living cell is to segregate chromosomes between the two daughter cells [1]. The mother cell achieves this by forming a *spindle*: a microscopic structure made of microtubules (MT) and cross-linking proteins, stretching across its length between the poles [2]. Once the spindle is formed and the chromosomes are aligned at the equatorial plane, the sister chromatids are pulled apart towards the opposite poles [1]. Chromosome segregation is known to play an important role in cancer progression, adaptation, and drug resistance [3–8]. Furthermore, a cell has to form a spindle within a suitable time-frame as a part of the regular cell-cycle. How a large number of signaling molecules, MTs, molecular motors, and the chromosomes self-assemble to form a spindle is a fascinating problem and has been a topic of active research [2, 9, 10]. One important aspect of spindle morphogenesis is the attachment of the MTs to the kinetochores (KC) [11] which are protein complexes on the chromosomes.

How does a MT attach to a KC? This question has been raised and studied before, and the mechanism by which this attachment takes place can vary across cell types [12–16]. One such mechanism for capture is based on the dynamic instability of MTs [17, 18]. Also known as *search-and-capture* (*S&C*), in this mechanism for capturing a KC, a large number of dynamic MTs grow from the centrosome in different directions [19, 20]. The KC attaches to one of the MTs when they both hit each other. *S&C* has been the most widely reported mechanism and the capture timescales for this mechanism has been well studied, both using a simplified theoretical approach [20–23], as well as using detailed computational models [15, 22, 24]. Recent *in vivo* experiments however suggest that simple *S&C* may not be the only way to capture the chromosomes. Existence of more sophisticated capture mechanisms have been reported. Some examples are: (a) biased MT dynamics due to Ran-GTP gradient around the KC [14, 22, 25, 26], (b) nucleation of MTs either from the spindle microtubules (also called *branching*) [27–31], or from the chromosome [15, 32–35]. Furthermore it has been observed that the motion of a KC and adaptive changes in its size may also affect the capture process [15, 16, 36, 37].

All these observed mechanisms involve some active chemical processes. It would be quite surprising if the goal can be achieved by a passive mechanism using lesser resources. A recent experiment by Kalinina *et al*. [16] reported such an interesting exception in *Schizosaccharomyces pombe*, commonly known as *fission yeast* — the KCs, which diffuse freely in the nucleoplasm, are captured by MTs which unlike spindle MTs in other organisms have a large duration of *pause* interrupting their usual dynamic instability. During the paused state, a MT is not stationary but executes *rotational diffusion* [16, 38, 39] (henceforth we will denote it as *RD*) being pivoted at the spindle pole body (SPB), where its random angular movement is driven by the thermal noise of the surrounding nucleoplasm. It is important to note that unlike other organisms, where the MT to KC ratio is quite large (for example, in human cells, ~ 17 MTs per KC [11, 40]), in *fission yeast*, fewer spindle MTs are available to capture a KC (~ 3 to 5 MTs per KC) [16, 41]. A natural question arises: is there any relationship between the mechanism of KC capture and the number of spindle MTs available for capture in an organism? Addressing this question would require a systematic study of capture times for different mechanisms, as a function of MT number *N*. To the best of our knowledge, such a *comparative study* with varying *N* and for a *moving* KC has not been studied earlier in the literature. Study of capture time dependence on *N* has been done for *S&C* mechanism [22], and a heuristic functional dependence was suggested based on the approximation of static KC (see Sec. II for details). Recently, in a different context of MTs searching a static immunological synapse in T cells [42], a *N* dependence of mean capture time was studied.

The mechanisms of spindle assembly in *fission yeast* have been studied in several recent works [39, 43–45]. Focussing on the simpler question of KC capture, the observed *RD* mechanism [16], also referred to as “sweeping” motion of MTs in an earlier work [13], may seem to be exceptional due to some specific reasons. Unlike other organisms, in *fission yeast* the mitotic spindle-size is much smaller [46, 47] and is formed inside the nucleus [13, 48]. Also, the cold treatment done to the cells [16] might have affected the length distribution of MTs. The *RD* mechanism was shown to be robust to temperature variations — similar results were reported for 14°C, 24°C and 32°C [16]. On the other hand the experiments [13] and [24] (done at 37°C) reported faster kinetics. While for any mechanism, physical parameters like the confining volume, MT size distribution, temperature, would affect the capture process, yet, one would be curious to identify the most distinguishing characteristics of one mechanism from another which go beyond parameter variations. A computational study was done for a single MT [24] and it concluded that angular diffusion indeed impart advantage in capture when combined with dynamic instability, but by a meagre amount of ~ 25%. Another study on KC capture during meiosis compared different mechanisms of capture for a fixed number of MTs and KCs [38]. We note that the cases of biological interest whether it be *fission yeast* or higher eukaryotes all have multiple MTs (*N* > 1). In this work, we computationally study the statistics of *first capture times* [49, 50] of a KC by varying *N* systematically, and compare the different scenarios — our aim is to unravel the advantages and drawbacks of *RD* over *S&C*.

One major finding from this work is that while there is a temporal advantage of *RD* over *S&C* for fewer MTs, with increasing *N* that ceases to be the case. This role-reversal is not apparent by looking at the *mean* capture timescales, a measure that is commonly used to characterize a stochastic capture event. One needs to be careful, as the sampled timescales of the first capture in confined geometries often do not exhibit a central tendency — hence the *mean* time is inadequate and often a misleading representation of the stochastic times [51–54]. We highlight the weakness of this measure and argue for a more robust measure, namely the *typical* time. *Typical* time is associated with the tail of the capture-time distribution. Although this timescale represents capture events with low occurrence probabilities, yet, should be of interest to experimentalists since they set the upper limit to the timescales for completion of an event. We adapt an algorithm [55, 56] to compute the fraction of lost KC to a strikingly high precision [~ 10^−17^], and thus obtain the typical times with high accuracy. We further propose a new alternate method to estimate typical time by using *extreme value statistics* [57, 58], which may be useful to experimentalists. A careful study of the typical time shows that for large *N*, the mechanism of *S&C* is decisively advantageous over *RD*, rationalizing its ubiquitous presence in most eukaryotic cells.

## I. RESULTS

We numerically study the first capture of a KC to one of the *N* MTs inside the *fission yeast* nucleus (see Fig. 1(A)). Previous *in vivo* experiments [16] found that the MTs exhibit a state of pause with *RD*, which lasts for ≈ 2.05 min, between the usual growth state (of ≈ 0.56 min) and shrinkage state (of ≈ 0.39 min), while the KC diffuses inside the nucleoplasm (see Fig. 1(B)). Experiments showed that the KC can attach to the MT tip (*tip capture*) or, at any point along the length of the MT body (*lateral capture*) (see Fig. 1(C)). In the following sections, we present the results for *lateral capture* while the results for *tip capture* are discussed in the Supplementary Information (*SI*). Here, our primary aim is to compare the efficiency of different capture mechanisms as a function of MT number *N*. This is done by comparing models that allow us to separately identify the relative contributions due to chemical kinetics and that due to mechanics. First, we study a model (already introduced in [16]) in which growth-shrinkage kinetics of the MT is ignored. Preformed MTs of typical length *L_MT_*, pivoted at the SPB, sweep the space in search of a KC — this case of pure rotational diffusion of MTs will be referred as the *RD* model (see Fig. 1(D)). Similarly, one may imagine a hypothetical scenario in which the MTs in *fission yeast* did not have a pause state, like in many other eukaryotes, but a switch directly from the growth to a shrinkage state — we will refer to this pure search-and-capture case simply as the *S&C* model (Fig. 1(D)). We further compare these two cases with a third scenario in which there is growth-pause-shrinkage kinetics as seen in the experiments, but during the paused state, the angular motion of MTs are suppressed— this will be referred to as the *S&C+P* model (Fig. 1(D)). Although rather artificial, a similar *S&C+P* model has been considered earlier in a computational work but only for *N* = 1 [24]. Here we study the case for completeness, with varying *N*. Finally, to have a benchmark, we also study the full model that incorporates all the features observed in experiments, i.e. a MT performing both *RD* and growth-pause-shrinkage kinetics simultaneously — we will refer to it as the *RD+S&C* model henceforth (Fig 1(B) and Fig 1(D)). The simulation details of these four models are discussed in Sec. III. The parameters for all these models are taken from the *in vivo* measurements reported in [16] (See Table III for details).

**FIG. 1.**
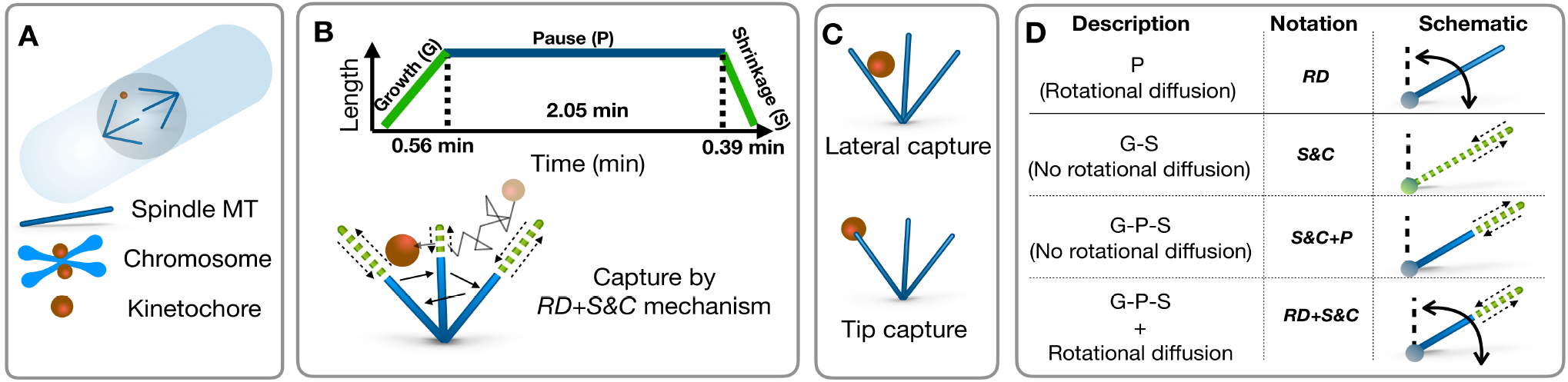
Study of KC capture in *fission yeast*. (A) Schematic of a *Schizosaccharomyces pombe* (*fission yeast*) cell. Unlike other organisms, here, the spindle is formed inside the nucleus. (B) Main features of KC capture in *fission yeast* are highlighted here. Length of a spindle MT is shown as a function of time (top). The average lifetime of a spindle MT is 3 min. There is a state of “pause” in between growth and shrinkage, where the MT length remains same. The MTs stay in this stationary state 70% of their lifetime and perform angular diffusion being pivoted at the SPB. The schematic (below) shows the growth, shrinkage and intermediate pause state with rotational diffusion, leading to the eventual capture of a KC. (C) A KC can attach either anywhere along the length or to the tip of the MT — these are referred to as *lateral* and *tip capture*, respectively. (D) The different models studied in this paper are schematically represented here. Blue solid color represents the pause state. Green dashed color represents the growth and shrinkage states. Straight and curved arrows represent the linear growth/shrinkage and angular diffusion, respectively.

The random times at which the capture of a KC happens by any one of the *N* MTs have a probability distribution function *F*(*t*), which is called the *first passage time distribution* [49]. The fraction of lost KC as a function of time which was experimentally measured [16], is referred to as the *survival probability* in the stochastic process literature and will be denoted by *S*(*t*) [49] — it is known that 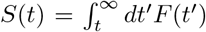. For capture processes in free space, *S*(*t*) typically has power-law tails (*S*(*t*) ~ *t*^−*γ*^, for *t* → ∞), while in confined geometries (like in our study, where the KC capture takes place inside the nuclear volume), it is expected to have exponential tails asymptotically: *S*(*t*) ~ exp(−*t/τ*) [49, 59]. Usually, the asymptotic behavior is hard to determine as obtaining *S*(*t*) to high precision is challenging. In this study using an algorithm of repeated enrichment [55, 56] (see Method 1 in Sec. III for details), we have obtained *S*(*t*) for the different models upto precisions ~ 10^−17^ (see Fig. 2(A)-(B)). One should note that multiple timescales are often present in the *S*(*t*) curves, which are not single exponentials (this is apparent in Fig. 2(A)-(B)) — this important feature has not been highlighted in earlier literature (see also Fig. S1 in *SI*). Moreover, all previous theoretical studies have focussed only on the *mean* capture times. In what follows, we would first discuss the mean capture time defined as 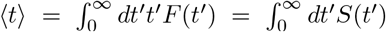. Then we would point out its limitations, and focus on the *typical* time *τ*, associated with the exponential tail of the *S*(*t*) function. We would argue that *τ* is a better representative than 〈*t*〉 to compare the *RD* and *S&C* mechanisms.

**FIG. 2.**
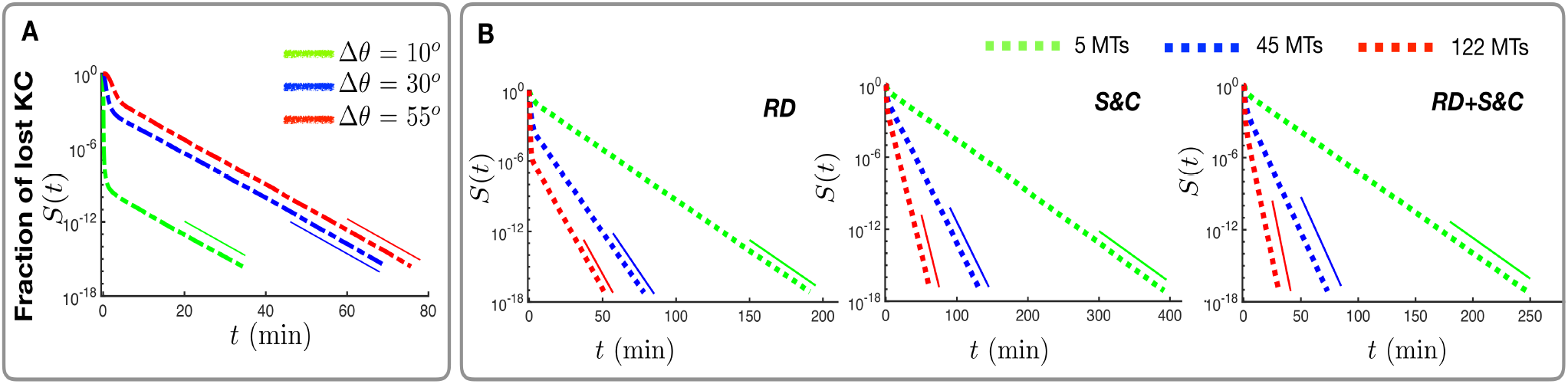
The fraction of lost KCs or survival probability *S*(*t*) as a function of time is a non-exponential function with an asymptotic exponential tail. (A) *S*(*t*) for *RD* mechanism with *N* = 45 MTs (all having same *θ_MT_*, but random *ϕ_MT_* initially) is plotted against time in a semi-log plot. Curves with different colors indicate different relative initial orientations (Δ*θ* = (*θ_MT_ − θ_KC_*)) of the MTs with respect to the KC (*θ_KC_* = 0 at *t* = 0). It is apparent that *S*(*t*) curves for three different initial conditions are all distinct. Yet at large times, the memory of the initial conditions go away — the *tails* of the curves become parallel to each other following exponential forms with the same decay constant. This shows that although mean times 〈*t*〉 are initial condition dependent, typical times *τ* are not. (B) *S*(*t*) curves for three different mechanisms are shown. Three different cases of *N* = 5,45,122 are shown in green, blue and red colors, respectively. As may be seen, the timescales of models *RD* and *RD+S&C* are close, while that of *S&C* are comparatively larger. The curves are non-exponential functions indicating the presence of multiple timescales, particularly for large values of *N*.

### A. Mean time of KC capture

Kalinina *et al*. showed that the dependence of the fraction of lost KCs on time can be nicely explained using the *RD* model for *N* ~ 3 – 5 MTs (see Fig. 3(c) in [16]). We estimate the mean capture time 〈*t*〉 ≈ 4.30 min from their experimental data (Fig. 1(b) in [16]). In Table I we have listed the various 〈*t*〉 for 5 MTs, computed using the different models (see Fig.1(D)). It is assuring that the *RD+S&C* model, which incorporates all the features in the experiments gets reasonably close to the experimental estimate (〈*t*〉_*RD+S&C*_ ≃ 6.52 min). Interestingly, for the *RD* model, 〈*t*〉_*RD*_ ≃ 5.38 min — it is almost same as the *RD+S&C* case, and in fact is marginally closer to the experimental value. This hints that if the MTs suffer catastrophe (which happens in the *RD+S&C* model but not in the *RD* model), a delay is introduced in the capture process. The fact that this *delay* is indeed due to dynamic instability, is further confirmed by study of the pure *S&C* model which has 〈*t*〉 almost twice compared to the *RD* model (see Table I). Finally, for the dynamically unstable MTs along with a pause state which do not rotationally diffuse (i.e. the *S&C+P* model), the time taken is even larger. This trend of ascending order of timescales with the elimination of angular diffusion continues even at higher *N* (see Fig. 3 and Tables S1 and S2 in *SI*). In particular, for the range of small *N* relevant to *fission yeast*, capture times are thus decisively minimised by the *RD* mechanism. On the other hand in *S&C*, time on an average is wasted due to repeated catastrophes — this is not a preferable strategy, particularly for organisms like *fission yeast*, having fewer spindle MTs. The 〈*t*〉 for all the models converge to the limit of mean MT growth time of ≈ 0.56 min at large *N*. Are there interesting statistical differences between the models at large *N* which is masked by this saturation in 〈*t*〉? Why is *S&C* so ubiquitous in the eukaryotic world?

**FIG. 3.**
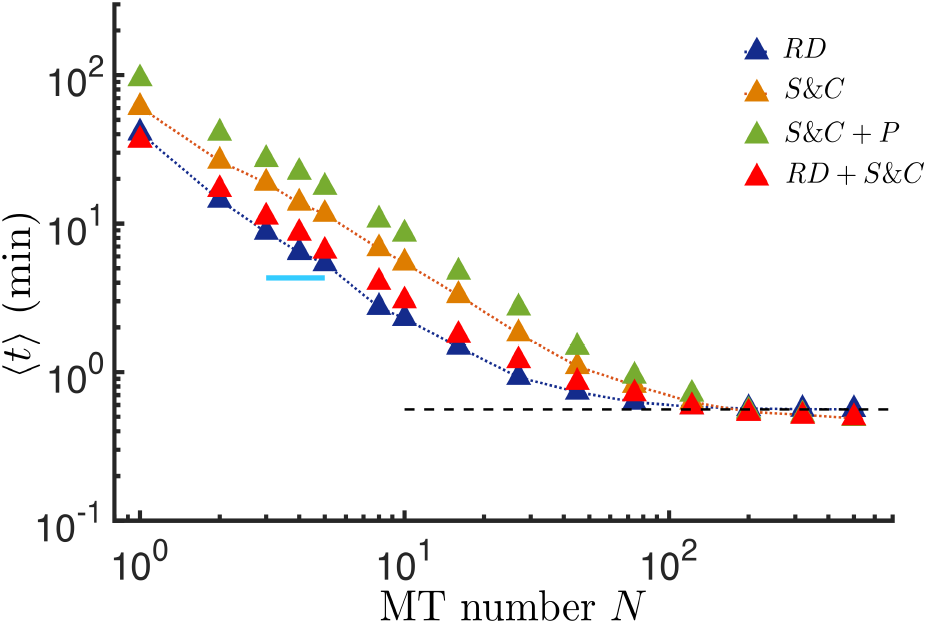
Comparison of capture-time efficiency using the mean capture time 〈*t*〉. We plot here 〈*t*〉 versus *N* on a log-log scale for all the cases shown in Fig. 1(D). *N* is varied from 1 to 500. Here we particularly highlight the important cases *RD* (blue) and *S&C* (orange) by plotting them with both symbols and lines. We also show the cases for *S&C+P* (green) and *RD+S&C* (red). Clearly *RD* is faster than *S&C*, as well as the other cases. Mean time of *RD+S&C* and *RD* are close over the full range of *N* studied. Mean time of *RD* for *N* = 5 is close to the experimental value (cyan line). At large *N* all the 〈*t*〉 values saturate to a threshold ≈ 0.56 min which is time taken on an average by the MTs to grow from the SPB to its average length *L_MT_*. Since the *RD* model does not take the initial growth of the MTs into account, we have added a growth time 0.56 min to 〈*t*〉_*RD*_ here, for comparison with other models.

**TABLE I.** Lateral capture times by 5 MTs

| Models | $\langle t \rangle$ (min) | $\tau$ (min) |
| --- | --- | --- |
| *RD* * | 5.38 * | 8.75 |
| *RD+S&C* | 6.52 | 10.87 |
| *S&C* | 11.56 | 16.39 |
| *S&C+P* | 17.58 | 22.22 |

\* $\langle t \rangle_{RD}$ value shown here includes a growth time value of 0.56 min. See Fig. 3 for details.

#### Is mean time an adequate representative of the KC capture times?

Mean time is very frequently used to characterize first passage processes in both theoretical and experimental studies, yet could be misleading due to two reasons. Firstly, the entire *S*(*t*) curve strongly depends on the initial spatial distribution of the KC and MTs. In our simulations, with the *RD* mechanism, we have chosen the *N* MTs to be initially oriented in random directions. This we believe is a generic initial condition. But in Fig. 2(A) we show that, if instead, the MTs were initially oriented in other special directions, the *S*(*t*) curves could be different. Consequently, the corresponding mean times would be distinct for the different initial conditions. Thus the mean first capture time is not a robust quantity to characterize the KC capture times, and is dependent on the initial arrangements of cellular components. This fact is not specific to this problem and well known in the stochastic process literature [49].

Secondly, we see in Fig. 2(B), that for small *N*, the *S*(*t*) curves are almost single exponentials, while with increasing *N*, existence of multiple timescales are clearly visible (see Fig. S1 in *SI*). In such cases, where the statistical data do not exhibit a central tendency, a mean value maybe misleading. Recent works on first passage problems in confined geometries have made such observations [52–54]. Any large variation of capture times between trajectories can be quantified by an indicator called the *uniformity index ω* [54]. If *t*_1_ and *t*_2_ are two instants out of a random set of capture times, then one defines *ω* = *t*_1_ /(*t*_1_ + *t*_2_). When times are not too different, *t*_1_ will be close to *t*_2_, and hence *ω* will be typically around 0.5. To the contrary, if *t*_1_ and *t*_2_ are very different, *ω* will be either close to 0 or 1. Thus by studying its distribution namely *P*(*ω*), we may conclude how non-uniform the timescales of capture are. By symmetry, swapping *t*_1_ and *t*_2_ should keep *P*(*ω*) invariant, i.e. *P*(*ω*) = *P*(1 − *ω*).

Since we wish to extract the real extent of trajectory to trajectory variations for each model, we subtract a common offset from the actual capture times, namely the minimum growth times of MTs before capture. From these set of modified capture times, we construct the probability distribution *P*(*ω*) as defined above. The results for the case of *RD* and *S&C* mechanisms, for different MT numbers *N*, are shown in Fig. 4(A) and Fig. 4(B), respectively. For the *RD* model (Fig. 4(A)), *P*(*ω*) has a peak near *ω* = 0 and 1, for both small as well as large *N*. Thus KC capture by this mechanism has large time variations across trajectories (for any *N*), and hence a mean value would not represent the frequently sampled times. On the other hand in the *S&C* case (Fig. 4(B)), while for small values of *N* there is a peak near *ω* = 0 and 1, with increasing *N*, the peak shifts near *ω* = 0.5. Thus for large *N*, mean values should be good representatives of the KC capture times by the *S&C* mechanism.

**FIG. 4.**
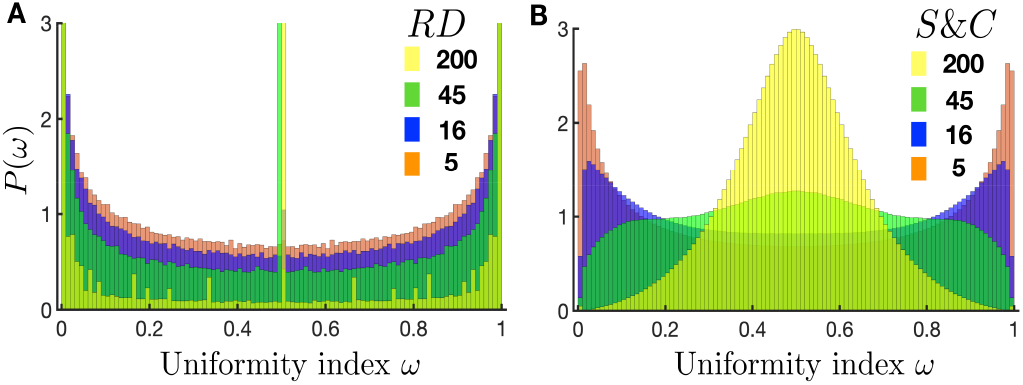
The adequacy of 〈*t*〉 is tested by studying the normalized distribution of the uniformity index *ω* (see Sec. IA) for different MT numbers (shown in different colors). For any given case, a deviation of *P*(*ω*) from a unimodal shape would arise due to large variability in the measured capture times. Here we show *P*(*ω*) for (A) *RD* and (B) *S&C* models. Clearly *P*(*ω*)|_*RD*_ is bimodal for all *N*, thereby showing that 〈*t*〉_*RD*_ is not a good representative of the overall statistics. *P*(*ω*)|_*S&C*_ on the other hand, is bimodal for small values of *N*, but becomes unimodal when *N* becomes large.

Given that *P*(*ω*) indicates large timescale fluctuations for small *N* (which is relevant for *fission yeast*), for both *RD* and *S&C*, we need a more robust measure to characterize the information of fluctuations contained in the processes. A natural thought would be to look at the *variance* of times. But we note that like mean times, variances would also be dependent on initial positions, since the whole *S*(*t*) curve does so (recall Fig 2(A)). Moreover, variances do not quantify the extreme temporal fluctuations. For that we need to study the *typical* capture time associated with the exponential tails of *S*(*t*), namely *τ*. For the rest of the paper, we shift our focus to this interesting quantity *τ*, which is also preferable as it is unaffected by any variation in initial conditions. We discuss this in details below.

### B. Typical time of KC capture

In Fig. 2(A), we see that although *S*(*t*) curves are distinct for different initial conditions, all the curves have parallel slopes at large time, indicating that they have the same decay constant *τ*. Thus the *typical* time *τ*, although dependent on MT number *N*, is independent of the initial spatial orientations of the MTs and hence is a rather robust quantity. Physically this happens as *τ* represents timescales of rare events of KC capture involving long trajectories of motion which remove initial memories. Biologically, extreme times are important to estimate upper bounds on completion of the KC capture process which affects mitosis. Since *S*(*t*) may be obtained rather precisely by a computational method (see Method 1 in Sec III), we obtain *τ* very reliably.

To compare the efficiency of the two mechanisms, we plot *τ* as a function of *N* in Fig. 5. For completeness, we compare the *τ* for all the four models (see Fig. 1(D)), but our primary focus is to highlight the differences between *RD* and *S&C* (plotted with symbols and dotted lines in Fig. 5). Importantly, when *N* is small, we see that *τ_RD_* < *τ_S&C_*. Thus for few MTs, *RD* is more efficient than *S&C* — a conclusion similar to the one shown above in Fig. 3 based on the comparison of 〈*t*〉, and consistent with the experimental findings of [16] for *fission yeast*. On the other hand, as *N* increases, *τ_RD_* starts saturating for 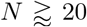 while *τ_S&C_* continues to decrease monotonically. This leads to a crossover between the two cases. Thus *S&C* mechanism is more advantageous temporally for cells with larger number of MTs. This is consistent with the observation that in many eukaryotes, where a large number (few hundreds) of spindle MTs are present [15, 40], dynamic instability driven capture of KCs are prevalent.

**FIG. 5.**
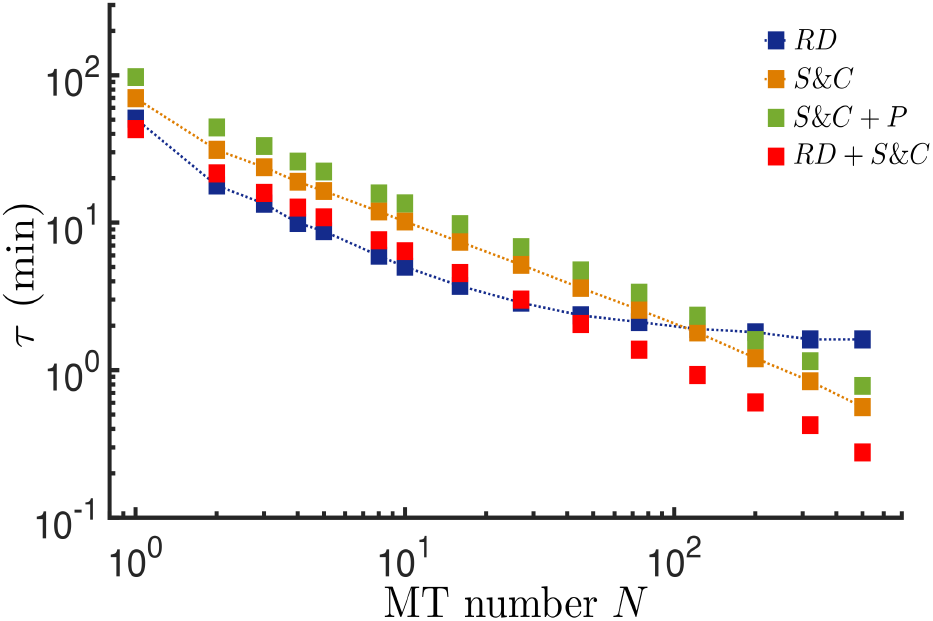
Comparison of capture-time efficiency by studying the typical capture time *τ*. Similar to Fig. 3, here *τ* versus *N* is plotted on a log-log scale for all the different cases (Fig. 1(D)). *N* is varied from 1 to 500. Again the main cases *RD* (blue) and *S&C* (orange) are highlighted with additional lines joining data points. For small *N* values, both *τ_RD_* and *τ_S&C_* decreases with growing *N*, with *RD* being more efficient than *S&C*. As *N* increases, *τ_RD_* starts saturating beyond *N* ≈ 20 while *S&C* keeps on decreasing leading to a crossover. The *τ* due to *S&C+P* (green) and *RD+S&C* (red) are also shown.

Fig. 5 also shows the typical times obtained from the *RD+S&C* and the *S&C+P* models respectively (plotted only with symbols in Fig. 5). Similar to Fig. 3, *τ_RD+S&C_* ≈ *τ_RD_* for small *N*. However as *N* increases, the contribution due to dynamic instability starts dominating leading to a crossover such that at large *N τ_RD+S&C_* < *τ_RD_*. This feature is also observed for the *S&C+P* model — although at small *N* it less efficient than the *RD* model (*τ_S&C+P_* > *τ_RD_*), as *N* increases, it becomes more efficient (*τ_S&C+P_* < *τ_RD_*).

#### Why does typical time *τ* saturate with increase in N for RD?

The saturation of the value of *τ* with increasing *N* in case of *RD* indicates that there exists a lower bound on time of KC capture that may be achieved by the mechanism. By adding more MTs beyond a certain number, the capture process cannot be made more efficient. What leads to such a saturation in *τ* (Fig. 5)? Note that, this saturation occurs at a higher value (≈ 1.56 min) than the saturation in 〈*t*〉 (≈ 0.56 min) (Fig. 3). In our model, the MTs have equal length, and therefore a MT tip diffuses on a curved surface which is a portion of a spherical surface with radius *L_MT_*, 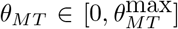 with 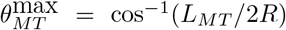, and *ϕ_MT_* ∈ [0, 2*π*]. There are two such surfaces accessed by MTs originating from both the SPBs. As *N* becomes large, the entire volume below (and above) the surfaces traced out by the tips of these numerous MTs (see Fig. 6(B)) becomes inaccessible to the KC since they get instantaneously captured there. To show that this is indeed the reason, we performed a separate simulation representing the *N* → ∞ limit — here the initial position of the KC is taken around the equatorial plane between two curved absorbing surfaces (see schematic in Fig. 6(B)). From this second simulation, we obtained a typical capture time 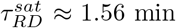. This is represented with a dashed line in Fig 6(C) — the re-plotted data of *τ_RD_* (already shown in Fig 5) completely converges to the dashed line at large *N*. This confirms our reasoning for the saturation based on geometrical constraint. The saturation of *τ* with increasing *N* in the *RD* model is a generic phenomenon due to the existence of a spatially absorbing continuous surface that restricts the KC diffusion within a sub-volume. To show this, we further present numerical and analytical calculations for a toy-model (see *SI*).

**FIG. 6.**
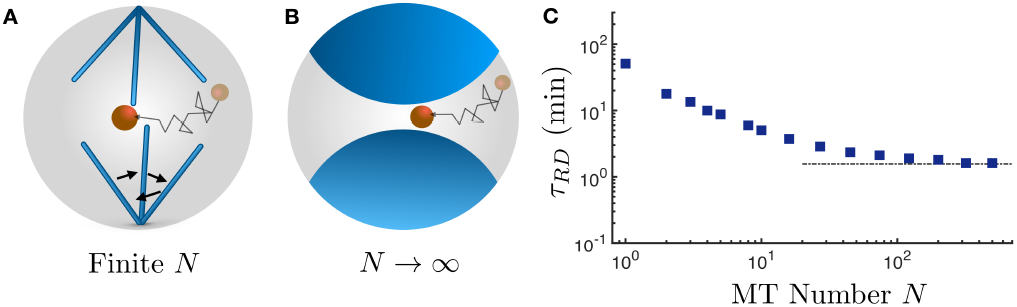
Explanation of the saturation of typical time *τ* in the *RD* model. (A) schematic of the *RD* model with finite number of MTs. (B) Schematic for *N* → ∞ limiting case. The *RD* model now consists of regions (sub-volumes shown in blue) which cannot be accessed any more. A KC starting from the central region can get captured at either of the surfaces bounding the sub-volumes. As a consequence in (C), *τ_RD_* saturates with growing *N* to a limiting value. This limiting value of 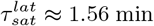, obtained by simulating the KC diffusion in presence of two curved absorbing surfaces (shown in (B)), is shown with a dashed line in (C).

### C. How to estimate typical time from experiments?

We have shown above how *typical* time *τ* is a very reliable quantity to compare the efficacy of different mechanisms of capture. While it is possible to estimate *τ* from the *S*(*t*) curves, which are obtained numerically up to high precision using Method I (see Sec. III), it may seem inaccessible to experimentalists who would typically have limited number of measured samples. For example, in [16], KC capture statistics was obtained using an assay with < 100 cells. Here we propose a method (see Method 2 in Sec III for details) based on *extreme value statistics* [57, 58], using which a reasonable estimate of *τ* may be obtained from limited available data. If the experiment consists of an assay with *N_t_ fission yeast* cells, each cell giving a random capture time, then the idea is to divide the measured times into *N*_2_ sets, each containing *N*_1_ = *N_t_*/*N*_2_ samples. The maximum times *t_max_* drawn from these *N*_2_ sets have a variance which is related to the typical time *τ* (see Eq. (4) in Sec. III). In Table II, we see that for *S&C*, for just *N*_1_ = 100 and *N*_2_ = 10, an experimentalist may get *τ* values not too different from the accurate values obtained by Method 1. For *RD*, we see in Table II that convergence to *τ* from Method 1 requires a relatively larger sample set, namely *N*_1_ = 1000 and *N*_2_ = 100. These results of *τ* from Method 2, just like the earlier results based on Method 1 discussed above, reconfirm that for small *N, RD* is more efficient, while for large *N, S&C* would be a preferable mechanism of capture. We hope that this Method 2 will be widely used henceforth by experimentalists, for studying the typical times in first passage processes in cellular biology.

**TABLE II.** Estimation of typical capture times for *RD* and *S&C*

| $N$ | $\tau_{RD}$ (min) | | | $\tau_{S\mathcal{E}C}$ (min) | | |
| --- | --- | --- | --- | --- | --- | --- |
| | Method-I | Method-II<br>$N_1 = 1000$ | | Method-I | Method-II<br>$N_1 = 100$ | |
| | | $N_2=10$ | $N_2=100$ | | $N_2=10$ | $N_2=100$ |
| 5 | 8.75 | 9.62 | 8.58 | 16.39 | 12.24 | 17.97 |
| 16 | 3.72 | 3.81 | 3.77 | 7.41 | 3.69 | 6.96 |
| 27 | 2.86 | 2.91 | 2.84 | 5.18 | 4.97 | 5.43 |
| 45 | 2.35 | 1.75 | 1.51 | 3.60 | 3.39 | 3.76 |

## II. DISCUSSION

Finding KCs by the spindle MTs is a crucial part of mitosis. The capture should happen fast enough to ensure uninterrupted progress of the cell division cycle. Due to both experimental and theoretical limitations, a complete physical picture of what dictates such a process is still lacking. The cell interior is a complex environment which is dynamic and crowded. Furthermore, the degree of this complexity can vary across organisms. Why do organisms often select different mechanisms to achieve capture? It is a moot question whether the existence of any specific mechanism is just a random selection, or, given the machinery and resources available to an organism, the chosen mechanism optimize something. A systematic study of first capture times as a function of MT number *N*, comparing different possible mechanisms is lacking. In this paper we address this issue in elaborate details, in the context of the biophysical problem of KC capture in *fission yeast*.

We started with the main question whether there is any advantage of choosing *RD*, which has only been reported in *fission yeast* [16, 38, 39], over the standard *S&C* mechanism. By numerically studying these two different mechanisms and by comparing their typical times *τ*, we have shown that *RD* is more efficient than *S&C* only when the MTs are fewer in number. This indicates that for any organism that has fewer spindle MTs like *fission yeast*, choosing *RD* over *S&C* can be advantageous in terms of time-optimality. Furthermore, the fact that this mechanism is thermally driven makes it advantageous on energetic considerations as well, than chemically driven *S&C*. We also find that for large number of MTs, the *S&C* mechanism becomes more efficient than angular diffusion — this may explain the common occurrence of the latter in eukaryotes.

The fact that *fission yeast* uses *RD* to search KCs instead of the other standard mechanisms has been a matter of surprise to cell biologists. It is possible that under special circumstances, *fission yeast* uses a combination of both *RD* and *S&C*. Such a mixed mechanism was studied computationally in [24] for a single MT and compared with the case where the MT undergoes dynamic instability with a paused state and no angular diffusion (similar to our *S&C+P* case). In this work, we have studied *N* > 1, and systematically compared mean and typical times (see Table. I, Fig. 3, Fig. 5 and Fig. S1, Table S1, Table S2 in *SI*) for all the mechanisms (Fig. 1(D)). The fact that *RD* gives least capture times stands out. There is a rather simple physical argument for this. For a few MTs, it is obvious that *RD* helps them explore a larger solid angular space than *S&C* in a fixed time. In the latter case, mis-oriented MTs and their times of growth-shrinkage may both lead to additional delays in the first encounter with the KC.

In this work, we have raised an important question: what is a robust measure to quantify the capture time statistics of the KCs? In all previous studies on KC capture, mean first-passage time 〈*t*〉 has been used to study the capture [15, 22, 23]. Mean times (undesirably) depend on initial conditions, and moreover the meaningfulness of 〈*t*〉 is often in question for broad distribution of capture times. To emphasize this point, we studied the distribution of the uniformity index *P*(*ω*), which if not unimodal at *ω* = 0.5, indicates huge variation across trajectories. We show that the case of *RD*, the fluctuations persist for all *N*, while in the case of *S&C* the fluctuations reduce as *N* becomes large. This can be understood in a heuristic way. The first passage distributions for both *RD* and *S&C* are not pure exponentials, having contributions of various short and long timescales. The long timescales arise from long diffusive excursions of a KC which escapes hitting any of the MTs. In case of *RD*, since KCs and MTs diffuse in the space available — although the mean time reduces with increasing *N*, the diversity of capture times due to very long and very short excursions persist for any *N* (Fig 4(A)). On the other hand, for *S&C*, since MTs move rectilinearly in 3d space, they appear as static line traps in the path of the diffusing KCs. Thus in this case, with increasing *N*, there is a fall in the number of long excursions due to a rise in the number of these traps, making *P*(*ω*) more unimodal (Fig 4(B)). This finding that at large *N*, 〈*t*〉 is a good measure for the *S&C* mechanism, is a good news for all the previous studies where the 〈*t*〉 has been measured for organisms with large number of MTs [15, 22].

We use typical time *τ* to distinguish between the capture events due to the two mechanisms with varying *N* (Fig 5). Although *τ* is a robust quantity, estimating it may seem challenging *a priori*, as it is associated with events of low probability. In this work, following an algorithm based on successive cloning of the system, we demonstrated that *S*(*t*) may be obtained to a high degree of precision (~ 10^−17^) (Fig 2). Thus a very accurate estimate of *τ* is possible in practice, and we hope that this method could be extensively used in future in computational studies of similar problems. Furthermore, we suggested an alternate method based on the theory of extreme value statistics, which may provide an approximate estimate of *τ* (see Table II, Method 2 in Sec. III). This may be of practical importance to experimentalists with access to limited sample sizes, nevertheless seeking an estimate of the upper bound of times for first capture.

We have shown that *τ* saturates with an increase in *N* when MTs execute *RD*. This saturation of *τ* is expected to be quite general and may be seen in other organisms with pivoted MTs — this is because pivoted MTs cannot explore more than a restricted hemispherical region of space even for *N* → ∞ (Fig 6). An implication of this saturation in *τ* with *N* for *fission yeast* is that, increased number of microtubules do not reduce the time of capture beyond a certain point. On the other hand for *S&C, τ* can in principle be reduced indefinitely as the entire available volume becomes accessible with increasing *N*. This implies that *S&C* will be preferred to *RD* by organisms having large number of spindle MTs. To show the crossover clearly, in our computational study, we considered *N* upto few hundreds. However, what is more important is the onset of saturation of *τ_RD_* around 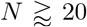, which is not very large.

How do the capture times scale with *N*? Earlier studies on *S&C* had suggested a heuristic form of 〈*t*〉 ~ 1/*N* based on certain simplified assumptions including a *static* KC [22]. For the more biologically relevant case with moving KC, this conclusion is no longer valid and requires a systematic study. Here we would like to summarize some of the facts which have emerged from our study. For a simple case with a static KC, the typical time *τ* ∝ 1/*N*, as in this case the problem simplifies to *N independent* capture events by *N* MTs [49]. On the other hand, the motions of the *N* MTs relative to the moving KC get *correlated*, thus leading to non-trivial dependence of the mean and typical times on *N*. For the case of *S&C*, the typical time *τ* shows a power law decay ~ *N*^−*β*^, with an exponent *β* ≈ 0.7 over a considerable range of *N* (see Fig. S4(A) in *SI*). For the mean time 〈*t*〉, a power-law dependence is seen only over a limited range of *N* with an exponent value ≈ 1.1 (see Fig. S4(B) in *SI*). Although here we numerically study the *S&C* process for the case of *fission yeast*, such *N*-dependence may be present in other capture problems under confinement. Interestingly there is a classic problem in stochastic process literature of *N* lions chasing a moving lamb, which is very similar to our problem of *N* MTs searching a KC. While some results are known in free space for that problem [60, 61], for a closed volume like a cell, our unpublished results [62] show diverse power-law dependences of *τ* on *N*.

In this work, for simplicity, we have ignored the finite-volume of the MTs. For a small number of MTs, as in the case for *fission yeast*, this assumption holds true. As *N* is increased, the excluded volume effects might become important. Moreover, there is an upper limit on the number of MTs that can be accommodated on the surface of a SPB (see *S’I*). For *S&C*, since MTs growing radially out of the SPB do not explore the volume laterally, this effect has been neglected as in earlier works [15, 22]. For the *RD* mechanism, this effect will become important beyond a certain MT number — this may be studied using more sophisticated computational models in future. However, the broad conclusions that we draw from this work, like the relative capture time efficiency of *RD* over *S&C* for different ranges of *N* are expected to remain unchanged. Moreover, the saturation that we see in *τ* with increasing *N* for *RD*, is expected to occur at a lower value of *N*.

It might be worth speculating, if there are further advantages of choosing *RD* over *S&C* in *fission yeast*. The *RD* and *S&C* mechanisms may be compared through two temperature (*T*) dependent parameters: diffusion coefficient (*D*) and reaction rate(s) *k*. Since *D ~ T* and *k* ~ exp(−const/*T*) (Arrhenius law) [63], a small change in temperature would cause a large change in *k* in comparison to *D*. As a result a diffusion driven random process will be comparatively less sensitive to temperature variations. This fact might be beneficial for *fission yeast* to cope with large temperature variations.

We hope that our findings may encourage future studies comparing capture mechanisms in different organisms with varying spindle sizes (and hence varying number of MTs). This may test our basic hypothesis that physical advantages of one mechanism over others may drive their selection.

## III. MATERIALS AND METHODS SIMULATION METHOD OF THE MODELS

Here we first discuss the computational models to study the attachment of MTs to the KC by *RD* and *S&C*. The KC is assumed to be a small sphere of radius *a*, which diffuses freely in the nucleoplasm. The nucleus is modelled as a sphere of radius *R*. MTs can nucleate from either of the spindle-pole-bodies (SPB). All the parameter values used in our simulations (see Table III) were taken from experimental measurements done at *T* = 24° C [16].

### A. Kinetics of the Kinetochore

For all the four models that we study in this paper (Fig. 1(D)), the KC dynamics is identical. The overdamped dynamics of KC [16, 50], is given by

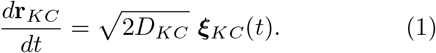

**TABLE III.** Parameter values used in simulation

| Models | Parameters | Values |
| --- | --- | --- |
| *RD* | Radius of nuclear sphere ($R$) | $1.55 \mu\text{m}$ |
| | Radius of KC ($a$) | $0.124 \mu\text{m}$ |
| | Length of MT ($L_{MT}$) | $1.5 \mu\text{m}$ |
| | $D_{KC}$ | $0.0354 \mu\text{m}^2\text{min}^{-1}$ |
| | $D_{MT}$ (angular diffusion coefficient) | $0.0603 \text{rad}^2\text{min}^{-1}$ |
| *Dynamic Instability* | $D_{KC}$ | $0.0354 \mu\text{m}^2 \text{min}^{-1}$ |
| | Tubulin subunit length | $6.15 \times 10^{-4} \mu\text{m}$ |
| | Growth rate ($r_+$) | $4387.5 \text{ subunit min}^{-1}$ |
| | Shrinkage rate ($r_-$) | $6175 \text{ subunit min}^{-1}$ |
| *S$\mathcal{E}$C* | **G** to **S** frequency ($f_{+-}$) | $1.8 \text{ min}^{-1}$ |
| | **S** to **G** frequency ($f_{-+}$) | $0 \text{ min}^{-1}$ |
| *S$\mathcal{E}$C+P* | **G** to **P** frequency ($f_{+0}$) | $1.8 \text{ min}^{-1}$ |
| | **P** to **S** frequency ($f_{0-}$) | $0.49 \text{ min}^{-1}$ |
| | **P** to **G** frequency ($f_{0+}$) | $0 \text{ min}^{-1}$ |
| | **S** to **P** frequency ($f_{-0}$) | $0 \text{ min}^{-1}$ |
| *RD+S$\mathcal{E}$C* | $D_0$ | $0.2036 \text{ rad}^2 \text{min}^{-1} \mu\text{m}^3$ |
| | $\tilde{D}_{MT}$ | $D_0/r_{MT}^3$<br>(if $r_{MT} \geq 1.5 \mu\text{m}$),<br>$0.0603 \text{ rad}^2\text{min}^{-1}$<br>(if $r_{MT} < 1.5 \mu\text{m}$) |
| | **G** to **P** frequency ($f_{+0}$) | $1.8 \text{ min}^{-1}$ |
| | **P** to **S** frequency ($f_{0-}$) | $0.49 \text{ min}^{-1}$ |
| | **P** to **G** frequency ($f_{0+}$) | $0 \text{ min}^{-1}$ |
| | **S** to **P** frequency ($f_{-0}$) | $0 \text{ min}^{-1}$ |

Here, **r**_*KC*_ is the position vector, and *D_KC_* is the translational diffusion coefficient of the KC. The components of ***ξ***_*KC*_(*t*) are Gaussian white noise with mean zero and delta correlations 〈*ξ_i_*(*t*)*ξ_j_*(*t*′)) = *δ_i,j_δ*(*t* − *t*′) with *i, j* = *x_KC_, y_KC_, z_KC_*. The initial position of the KC is uniformly distributed inside a small sub-sphere around the center of radius 0.3 *μ*m. At the nuclear boundary, reflecting boundary condition is applied along the radius vector joining the KC and the center of the nuclear sphere.

### B. Langevin simulation for Rotational Diffusion of MTs

In this process, all the MTs are assumed to have fixed lengths *L_MT_*, pivoted at either of the SPBs (see Fig. 1(D)). In the overdamped limit, the equation of motion for the MT in spherical polar coordinates is given by [16]

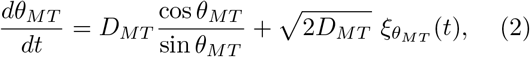

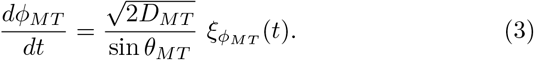

Here, *D_MT_* is the angular diffusion coefficient of the MTs. Similar to *ξ_KC_*, here *ξ_MT_* = (*ξ_θ_MT__, ξ_ϕ_MT__*) is a Gaussian white noise with mean zero and delta correlations. The initial orientation of the MTs pivoted at the SPBs are distributed uniformly inside the nuclear envelope. Since MT length *r_MT_* = *L_MT_* is a constant, the tip of a MT is always constrained to diffuse on a portion of a spherical surface of radius *L_MT_*, with 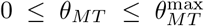 and 0 ≤ *ϕ_MT_* ≤ 2*π*. Here 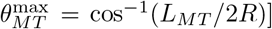. Reflecting boundary conditions are applied when the MT tip hits the boundary.

### C. Langevin-Gillespie hybrid algorithm for Search and Capture

The search-and-capture *S&C* mechanism involves dynamic instability of MTs [17, 20]. Experiments [16] however showed that the MTs exhibit a state of *pause* (**P**) before switching from the *growth* (**G**) to the *shrink* (**S**) state (also adapted in [24]). To compare the relative efficiency, here we study different models of *S&C* both *with* or *without* the **P** state. MTs have a tubular structure consisting of typically 13 proto-filaments [1]. Thus when 13 tubulin dimers are added (or subtracted), length of a MT *r_MT_* increases (or decreases) by a length of 8 nm. Since we do not have explicit protofilaments in our models, we take care of this by choosing the effective subunit length of each dimer to be 8/13 nm in simulations as has been done in earlier works [64–66]. In all our dynamic instability models, each MT starts out in a random direction with a ‘seed’ length of 20 dimers (i.e. 20 × 8/13 ≈ 12 nm) and grows by adding subunits with rate *r*_+_. In **S** state, a MT shrinks back to the cutoff length *l_min_* by losing subunits with rate *r*_−_. Subsequently, the MT switches back to the **G** state choosing a new random direction.

To simulate our models, we use a combination of Langevin dynamics and kinetic Monte Carlo. The KC position is updated using the Langevin equation (Eq. (1)), whereas the dynamic instability of a MT is modelled using Gillespie algorithm [67].

#### 1. Pure search-and-capture model (*S&C*)

This is the standard dynamic instability model where the **P** state is ignored (see Fig. 1(D)). MT switches from the **G** to the **S** state with catastrophe frequency *f*_+−_ keeping its orientation fixed. When a growing MT tip hits the nuclear envelope, it switches to the **S** state keeping its orientation unchanged.

#### 2. Search-and-capture with stationary MTs in pause state (*S&C+P*)

In this case a MT switches from **G** to **P** with frequency *f*_+0_ and **P** to **S** with frequency *f*_0−_ keeping its orientation fixed (see Fig. 1(D) and Table III). Upon hitting the nuclear envelope, the MTs switch to the **P** state.

#### 3. Search-and-capture with rotational diffusion (*S&C+RD*)

This model incorporates all the features observed in the experiments. A MT undergoes both rotational diffusion and dynamic instability (i.e **G→P→S**) simultaneously (see Fig. 1(B) and Fig. 1(D)). The state of MT remains unchanged on hitting the nuclear envelope. In this model unlike *S&C* or *S&C+P*, MTs not only have a Gillespie update due to growth, pause and shrinkage, but also has a simultaneous Langevin update due to rotational diffusion following Eqs. (2–3). But while the Langevin updates of KC and MT are more frequent and happen at chosen time intervals Δ*t*, the Gillespie updates for MT growth, pause and shrinkage events happen occasionally after several such Δ*t* intervals.

Since the length *r_MT_* varies, we chose a length dependent angular diffusion constant 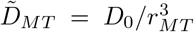 [68–70] for *r_MT_* > *L_MT_*, where 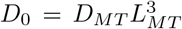 (see Table III). For *r_MT_* ≤ *L_MT_*, we use 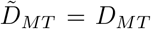 — note that having a 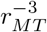 dependence all the way down to small values of *r_MT_* would lead to unrealistic values of 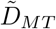.

### D. Capture Conditions

The positions of the MT tip and the center of mass of the KC are given by **r**_*MT*_ = (*r_MT_, θ_MT_, ϕ_MT_*) and **r**_*KC*_ = (*r_KC_, θ_KC_, ϕ_KC_*) respectively. The angle subtended at the SPB is:

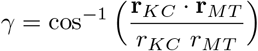

As discussed earlier, the KC can attach to the *tip* of the MT or *laterally* along the body (Fig.1(C)). The *lateral capture* condition is: *r_KC_* cos *γ* ≤ *r_MT_* and *r_KC_* sin *γ* ≤ *a*, or |**r**_*KC*_ − **r**_*MT*_| ≤ *a*. For *tip capture* (see Fig. S2 in *SI*), the capture conditions we use are in conformity with the criterion used by Kalinina *et al*. in their experiments [16].

## METHODS TO ESTIMATE TYPICAL CAPTURE TIME

For the randomly sampled first capture times of KC by MTs in the models discussed above, the typical times *τ* are challenging to estimate. We discuss below two different methods for estimating *τ*, which we have used in this work.

### E. Method 1: Cloning of systems to calculate *S*(*t*)

As discussed in Sec. IB, the asymptotic behavior of the survival probability *S*(*t*) is independent of the initial positions of the MTs and the KC and behaves as: lim_*t*→∞_ *S*(*t*) ~ exp(−*t/T*). However this *robust* asymptotic exponential tail may appear at very small values of *S*(*t*). Standard computational methods often fix a precision like ~ 10^−8^ for *S*(*t*) and try to get the tail behavior from large samples of *t*. Howsoever large number of samples are used, at such ordinary levels of precision, the asymptotic tail may not even appear and so accurate determination of *τ* remains a challenge. However an algorithm, developed earlier in the context of reaction-diffusion systems [55, 56], may be very effectively extended to study problems of survival in confined geometries and in particular get accurate estimates of asymptotic behaviour. The main idea of the algorithm is schematically depicted in Fig. 7. At *t* = 0, we start with *M* random realizations of the system — each one having *N* number of MTs and a KC. Initially the survival probability *S*(0) = 1. As time evolves, capture happens in some copies, while in the remaining (say *q*(*t*) copies the KC continues to survive. Thus at any time *t*, we have *S*(*t*) = *S*(0)(*q*(*t*)/*M*). At a time *t* = *t*_1_ when *q*(*t*_1_)/*M* = *s*_1_ just becomes ≤ 1/2 we replicate the *q*(*t*_1_) surviving copies to restore the initial ensemble size *M* — this step is referred to as *cloning* or *enrichment* [56]. Subsequently at any *t* > *t*_1_, if *q*(*t*) are the surviving copies, then *S*(*t*) = *s*_1_(*q*(*t*)/*M*) until the next enrichment event happens at *t* = *t*_2_ when *q*(*t*_2_)/*M* = *s*_2_ becomes just ≤ 1/2. This process is iterated many times. For *k* successive such cloning events *S*(*t*) = *s*_1_*s*_2_ ⋯ (*q*(*t*)/*M*) = *O*(1/2^*k*^), and thus accuracies like ~ 10^−17^ or even lower can be readily achieved. For such precision of *S*(*t*), it is guaranteed that the exponential tail clearly appears, and so *τ* is extremely accurately determined. In our simulations, the choice of a large number of realizations namely *M* = 1000 reduced fluctuations to a minimal, but the computational cost was fairly large. Each *S*(*t*) curve for a particular *N* of each model (Fig 1(D)), is generated by running the simulation for 96 hours, using the super-computing facility at IIT Bombay.

**FIG. 7.**
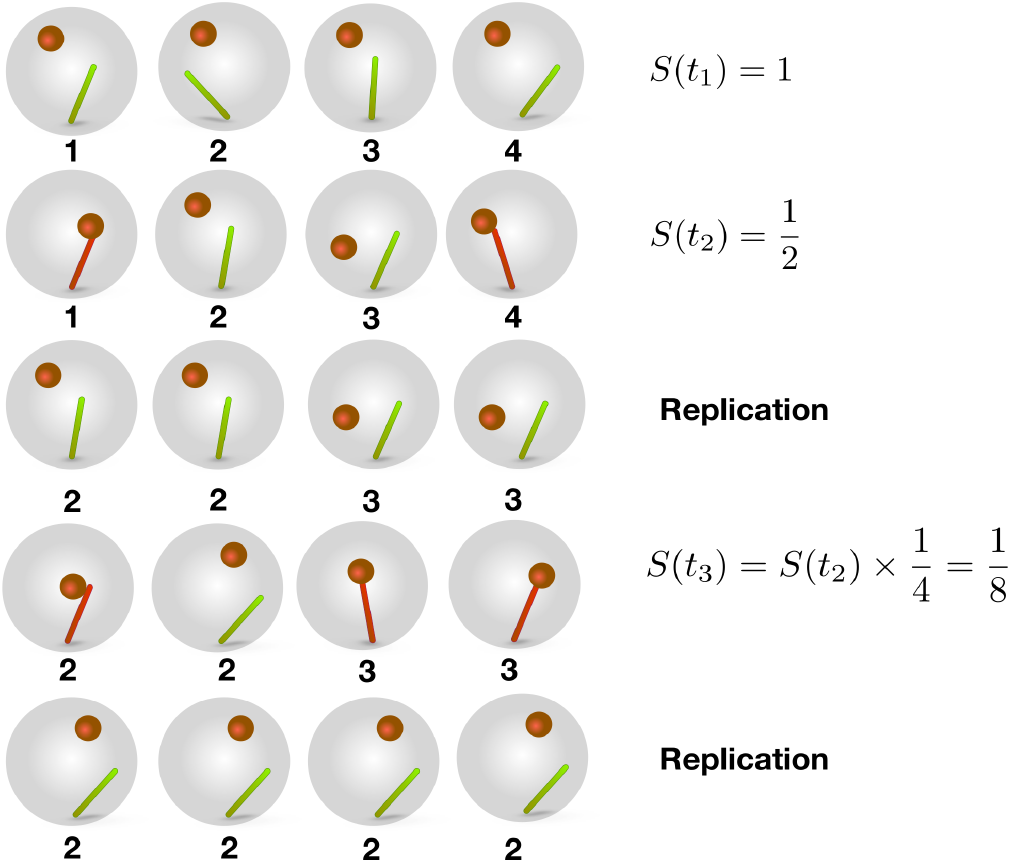
Illustration of the algorithm of Method-1 to calculate the survival probabilities *S*(*t*) to high precisions. In this figure, *M* = 4 and *N* =1.

### F. Method 2: Sampling extreme times

Extreme value statistics deals with extreme (maximum or minimum) deviations of a set of random observations. For a set of independent and identically distributed random variables {*t*_1_, *t*_2_,…, *t*_*N*_1__} that are drawn from a parent distribution *F*(*t*), one might be interested in distribution of the leader *t_max_* = max{*t*_1_, *t*_2_,…,*t*_*N*_1__}. It is known that if *F*(*t*) ~ exp(−*t/τ*) for large *t*, then the distribution of (*t_max_ − μ*)/*τ* approaches the universal Gumbel distribution [57, 58] for large *N*_1_. Note that *μ* and *τ* are non-universal constants dependent on the parent distribution *F*(*t*). Interestingly, the variance *σ*^2^ of the Gumbel distribution for *t_max_* is related to the decay constant of the exponential tail of *F*(*t*) as: *σ*^2^ = *π*^2^*τ*^2^/6 (see [58] and *SI*). Thus if one can experimentally or computationally estimate *σ*^2^, then one may invert this formula to get

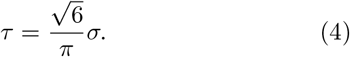

We propose that the above idea may be used for the purpose of estimating typical times of first capture of KC by MTs having the distribution *F*(*t*). Note that the decay constants of exponential tail of *F*(*t*) and survival probability *S*(*t*) are the same. In computation or experiment, one needs to first sample *N*_1_ × *N*_2_ different times of capture. The whole data may be divided into *N*_2_ sets, each having *N*_1_ sample times. Each such set may provide a *t_max_*, such that there would be *N*_2_ such values. These *t_max_* values may then be used to obtain *σ*^2^. Finally using Eq. (4) one may obtain *τ*. Table II in the text show the efficacy of this method.

## ACKNOWLEDGMENTS

A.N. acknowledges IRCC at IIT Bombay, India, and SERB, India (Project No. ECR/2016/001967) for financial support. D.D. and A.N. acknowledge MPIPKS, Dresden, for kind hospitality and computational facility. I.N. acknowledges the High Performance Computing Facility at IIT Bombay.

## Supplementary Material

### Ratio of typical and mean timescales — Indicator of non-exponential forms of *S*(*t*)

For a simple exponential function *S*(*t*) = exp(−*t*/*τ*), it is easy to see that the mean time 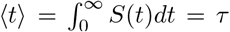 (typical time). But the curves *S*(*t*) which we obtain in our study (for example in Fig. 2 in the main text), have non-exponential functional forms. Nevertheless, they all have exponential tails. The presence of multiple timescales manifest in the fact that 〈*t*〉 = *τ*. Thus, the ratio of *τ*/〈*t*〉 is an indicator of the non-exponential form of *S*(*t*). In Fig. 1 below, we plot this ratio (by using the data of Fig. 3 and Fig. 5 in main text) as a function of *N* for the four different models studied in the paper. For a few MTs (< 3), the values of the ratios are slightly greater than 1, indicating that the corresponding *S*(*t*) functions are “almost” single exponentials. But with increasing *N* there is a strong departure from exponentiality indicated by the rise of the values of the ratios. In fact, for the RD model even at the largest *N* that we studied, the ratio stays very different from 1. This compliments our finding that *P*(*ω*) remains bimodal for the *RD* model for all values of *N* (hence showing large trajectory to trajectory fluctuations in capture times). For the other three models while 〈*t*〉 tends to saturate at large *N* (from Fig. 3 in the main text), the typical time *τ* continues to decrease (Fig. 5 in the main text). Consequently, the ratio *τ*/〈*t*〉 shows a plunge in Fig. 1 at large *N*.

**FIG. 1:**
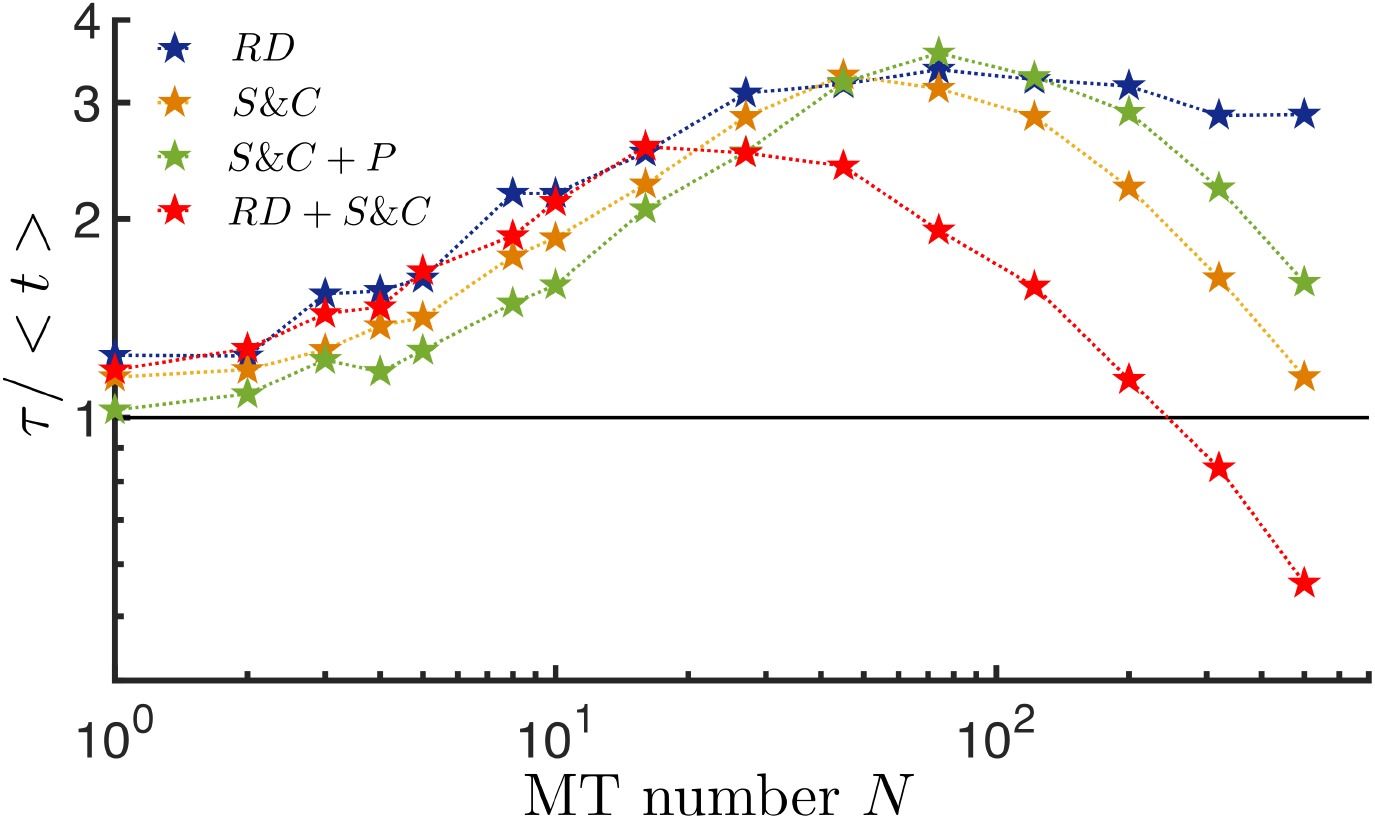
Ratio of the typical and mean capture times for the four models studied in this work as a function of MT number *N*. The black solid line indicates the case of simple exponential.

### Efficiency of the *RD* model with respect to the other models

As discussed in the main text in addition to the mean time 〈*t*〉, the typical time *τ* is a very useful quantity to study. Here, we compare the timescales of four models discussed in the main text to demonstrate the quantative temporal benefit that the *RD* mechanism produces. The efficiency of the *RD* model is expressed in terms of two ratios *τ_RD_*/ *τ_Model_* and 〈*t*〉_RD_/〈*t*〉_*Model*_ in the table below for MT number 1, 3 and 5, respectively. Amongst the three *N* values studied below, the efficiency of the pure *RD* mechanism is most clearly manifested for *N* = 5.

In Fig. 3 in the main text, we have shown that the 〈*t*〉_RD_ is the smallest compared to all the other models for a significant range of *N*. In Table II, we explicitly write some values of 〈*t*〉 for the four different models as a function of *N* in descending order of the temporal efficiency.

**TABLE I:** *τ_RD_* and 〈*t*〉_*RD*_ compared to other dynamic instability associated models.

| Model | $\tau_{RD} / \tau_{Model}$ | | | $\langle t \rangle_{RD} / \langle t \rangle_{Model}$ | | |
| --- | --- | --- | --- | --- | --- | --- |
| | $N = 1$ | $N = 3$ | $N = 5$ | $N = 1$ | $N = 3$ | $N = 5$ |
| $RD+S\mathcal{E}C$ | 1.18 | 0.84 | 0.80 | 1.10 | 0.74 | 0.74 |
| $S\mathcal{E}C$ | 0.73 | 0.57 | 0.53 | 0.66 | 0.44 | 0.42 |
| $S\mathcal{E}C+P$ | 0.52 | 0.41 | 0.39 | 0.43 | 0.30 | 0.27 |

**TABLE II:** 〈*t*〉 for different models as a function of *N*.

| Model | $\langle t \rangle$ (min) | | | |
| --- | --- | --- | --- | --- |
| | $N=5$ | $N=10$ | $N=45$ | $N=122$ |
| $RD$ | 5.38 | 2.29 | 0.73 | 0.58 |
| $RD+S\mathcal{E}C$ | 6.52 | 3.02 | 0.85 | 0.59 |
| $S\mathcal{E}C$ | 11.56 | 5.44 | 1.09 | 0.63 |
| $S\mathcal{E}C+P$ | 17.58 | 8.51 | 1.47 | 0.71 |

### Typical time for varying MT number for the case of tip capture

In the main text, we have discussed all the results in the context of *lateral* capture. Here, we study the typical time for the case of *tip* capture, as a function of the MT number. We study here the case, where the KC can attach anywhere within <0.5 *μ*m of the MT tip — this was the definition of tip capture used in the experiment [1]. We find that the qualitative features are similar to the lateral capture case (compare Fig. 5 in the main text with Fig 2 below). We see that the *RD* model is more efficient than the *S&C* model (and also the other two models) for smaller *N*. Similar to the case of lateral capture, here as well we find that *τ*_RD_ saturates at large *N*. All other *τ* values keep monotonically decreasing. The reason behind the saturation in *τ*_RD_ can be explained using the similar argument of geometrical constraint as in main text. Here, the *annular* volumes (between radii *L_MT_* and *L_MT_* – 0.5) on both sides of the nuclear sphere become the region of the capture as *N* → ∞. The condition for *tip capture* is: 0 ≤ (*r_MT_* – *r_KC_* cos *γ*) ≤ 0.5 and *r_KC_* sin *γ* ≤ *a*, or, |**r**_*KC*_ – **r**_*MT*_| ≤ *a*.

**FIG. 2:**
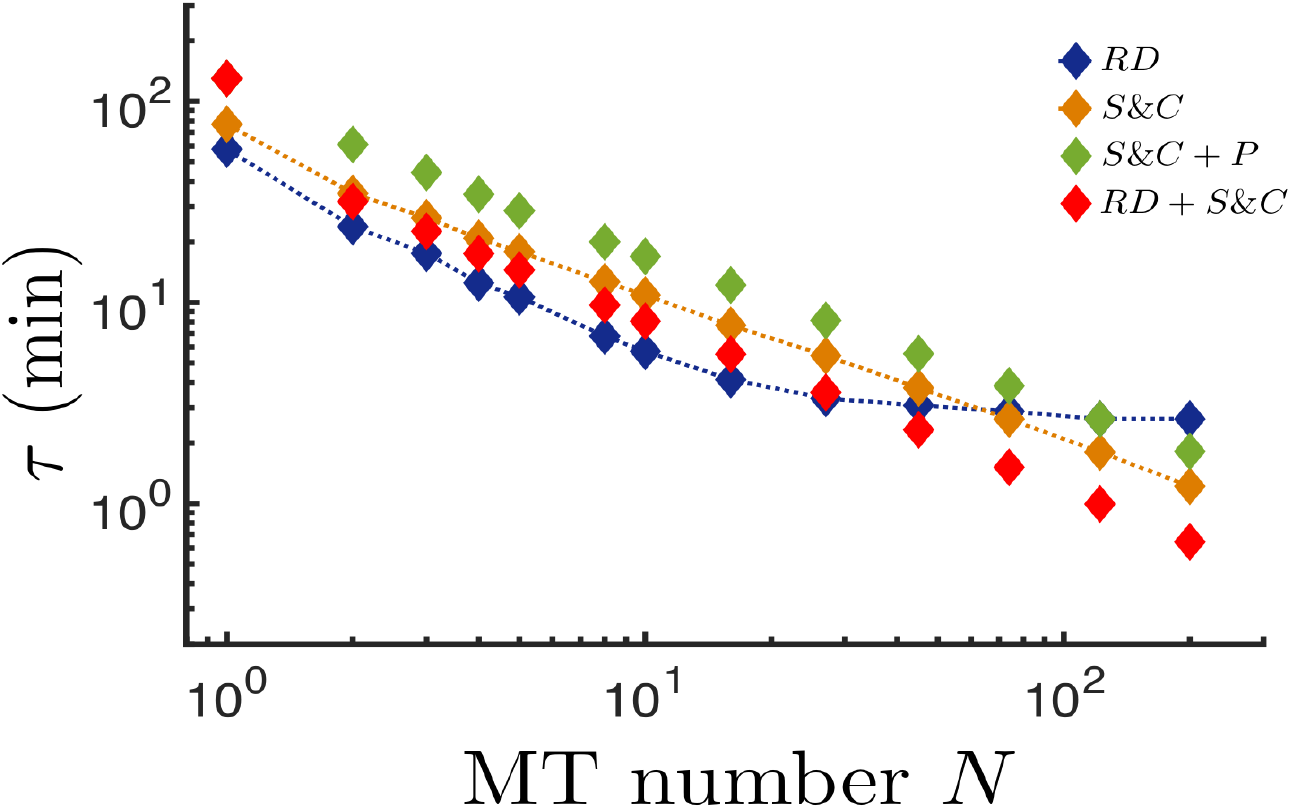
Typical times for tip capture as a function of *N*.

For large *N*, we have seen the saturation behavior of *τ*_*RD*_ in Fig. 6 in the main text and Fig. 2 above. Here we show that this behavior is generic by studying a toy-model analytically and computationally. Consider a circle of radius *b* inside which a small disc of radius *a* (like the KC) diffuses freely. *N* point particles (like the tips of *N* MTs) diffuse along the diameter (see Fig. 3(A)) of the circle. As *N* → ∞ the entire diameter becomes an absorbing line (see Fig. 3(B)). We perform simulations for finite *N* (see Fig. 3(A)) and plot the data in Fig. 3(C)). We have performed an analytical calculation for *τ* which yields *τ_sat_* ≈ 3.10 (a.u) — the latter is represented by a black solid line in Fig. 3(C). Convergence of the computational values of *τ* to the analytical value of *τ_sat_* for *N* > 10 is clearly visible. The analytical calculation is discussed below.

**FIG. 3:**
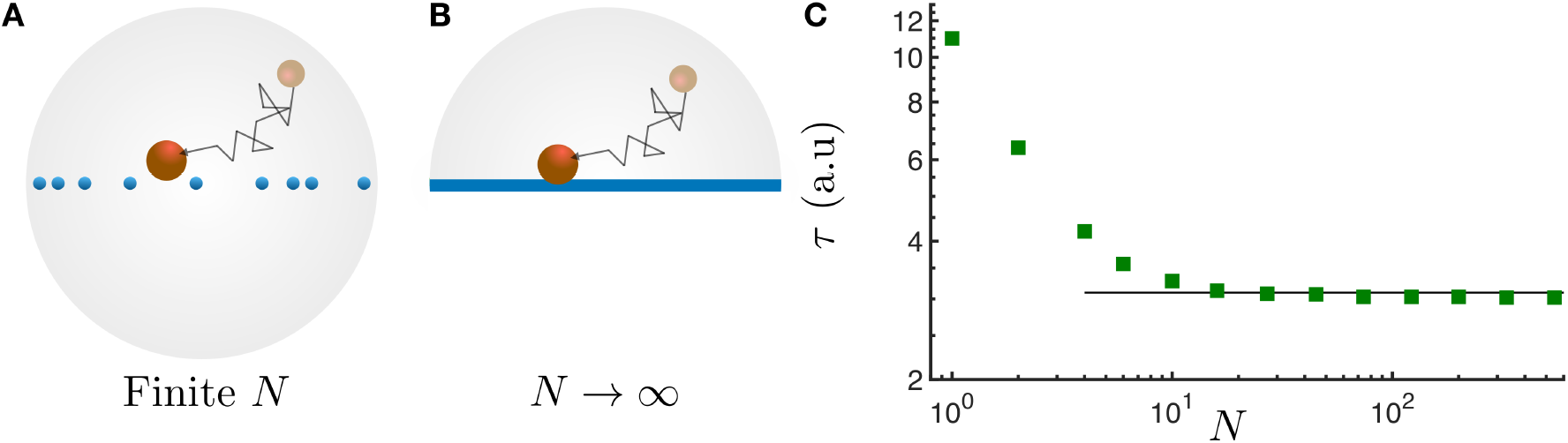
A toy model to explain the saturation is shown in (A). A disc diffuses inside a circle and can get captured along the diameter where non-interacting particles diffuse along the diameter. (B) The limiting case here is an absorbing line along the diameter of the circle which is realized for *N* → ∞. (C) The typical time **t** for the toy-model (green filled square) shows saturation with *N*. The *τ_sat_* ≈ 3.10 (a.u) (black line) is obtained from the analytical calculation discussed below.

### Analytical estimate of *τ_sat_* for the quasi-2d model

Using the *Backward Fokker-Planck* equation [2], we analytically estimate here the *τ_sat_* value for the quasi-2D model in the limit of *N* → ∞ (see Fig. 3(B)). The disk of radius *a* (similar to the KC) diffuses inside a semi-circle of radius *b*. The diameter of the semi-circle is an absorbing boundary while the semi-circular arc is a reflecting boundary. For the simplest case where disc is assumed to be as a point particle (*a* = 0), the backward Fokker-Planck equation for the survival probability *S*(*r*, *θ*, *t*) is given by,

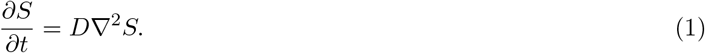

Where, ∇^2^ is the two-dimension Laplacian operator in polar coordinates. *S*(*r*, *θ*, *t*) is the survival probability of the particle upto time t starting from initial position (*r*, *θ*). Here *D* is the diffusion coefficient of the disc. We solve Eq. (1) using separation of variables. Substituting *S*(*r*, *θ*, *t*) = *R*(*r*)Θ(*θ*)*T*(*t*) in the above equation we get,

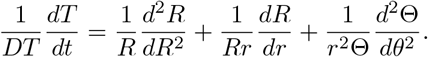

Equating the left hand side to a constant value −*k*^2^, we get the *T*(*t*) solution as

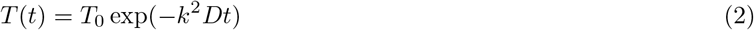

and the above equation becomes

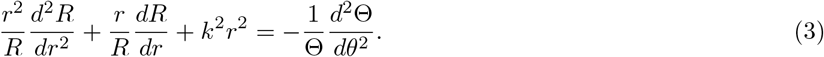

Equating right hand side to *m*^2^ and applying the absorbing boundary condition Θ(*θ*) = 0, at *θ*= 0 and *π*, we get the solution of the angular part as

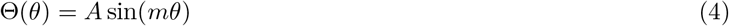

where, *m* can be any integer value. Finally, the radial part of the Eq. (3) gives the solution in terms of Bessel functions of order *m*,

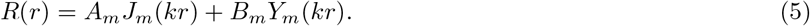

At *r* = 0, the radial function should vanish due to the absorbing boundary condition. Since *Y_m_*(*kr*) diverges as *r* → 0, we drop that term. The above relation is true for any positive integer *m*. The reflecting boundary condition leads to the vanishing of the first derivative of *R*(*r*) at *r* = *b*, i.e., 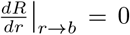. Using Eq. (5) and the reflecting boundary condition we obtain [3],

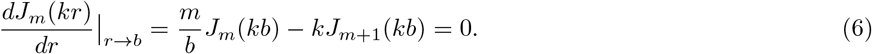

From Eq. (2) the typical time 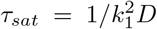, where *k*_1_ is the smallest positive value of *k* satisfying Eq. (6), corresponding to *m* =1. Assuming *k*_1_*b* is a small number, we approximate 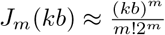 to finally obtain

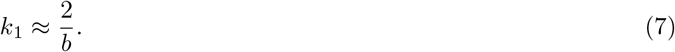

Since in our simulation the disc has a finite radius a (see Fig. 3(B)), and the capture happens as the periphery of the disc touches the diameter, we need to replace the radius *b* in Eq. (7) by an effective radius 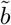. We use a geometrical appoximation 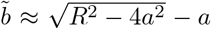. For *b* = 4.0, *a* = 0.4 and *D* = 1, we get 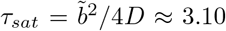 (a.u). We further confirm this value by directly simulating the case of Fig. 3(B) and find a numerical value of *τ_sat_* = 3.02 (a.u) which is in good agreement with the analytical approximation of 3.10 (a.u).

### Behavior of typical time *τ* and mean time 〈*t*〉 for *S&C* with *N*

In the main text, we mentioned the fact that *τ_S&C_* and 〈*t*〉_*S*&*C*_ have non-trivial dependence on *N*. To show that, here we replot the *τ_S&C_* and 〈*t*〉_*S&C*_ data (from Fig. 3 and Fig. 5 in the main text) as a function of *N*. As it is apperant, there is no clear power-law form spanning over the full range of *N*. We fit power-law form ~ *N*^−*β*^ over limited ranges of *N* in the two cases and extract the values of *β* (see Fig. 4). These turn out to be *β* = 0.7 and *β* = 1.1 for the typical and the mean times, respectively.

**FIG. 4:**
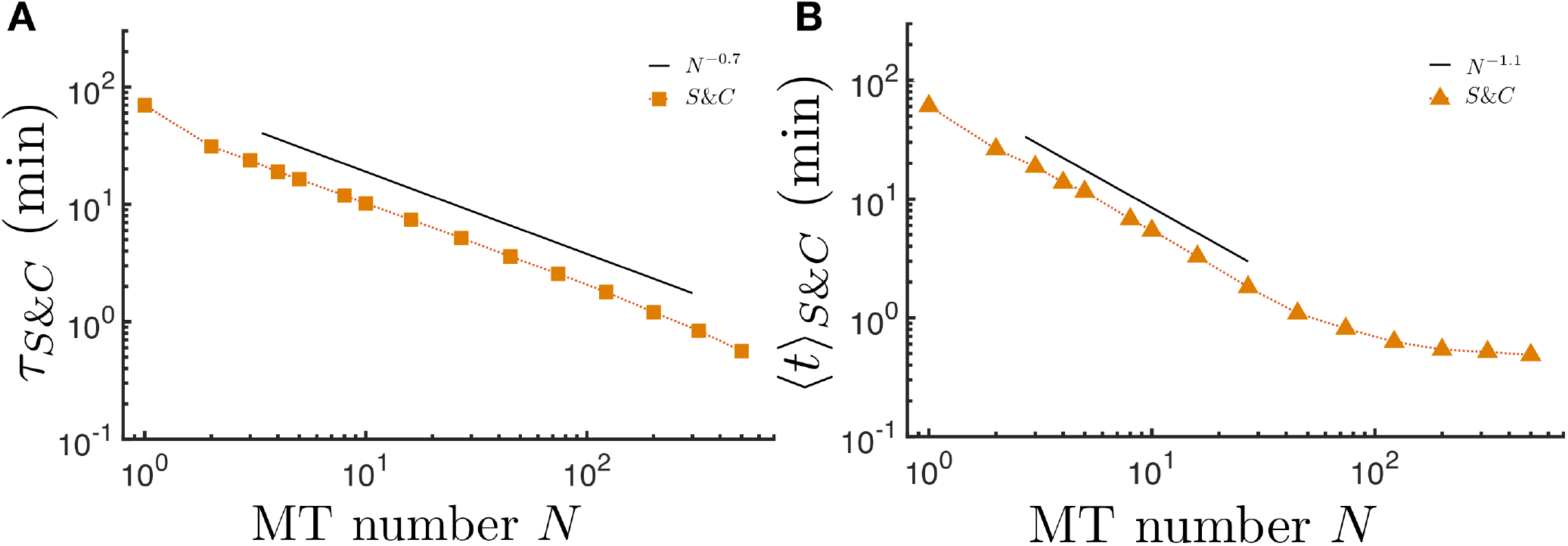
Power-law fits (black solid lines) over limited ranges of *N* of the data (from Fig. 3 and Fig. 5 in the main text) for (A) typical and (B) mean times for the *S&C* model.

### Estimation of maximum number MTs which may grow out of the surface of SPBs

The spindle pole body for *fission yeast* is oblate shaped with diameters 2*d*_1_= 0.18 *μ*m and 2*d*_2_ = 0.09*μ*m [4]. Assuming that MTs grow from the surface of SPBs, we may calculate the ratio of the surface area *S_A_* of a SPB and the cross-sectional area of a MT. This number gives an estimate of the maximum number of MTs that may grow out of a SPB. The area *S_A_* of a SPB is calculated by using the following formula [5],

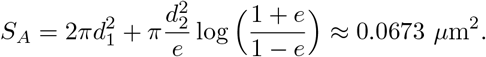

Here 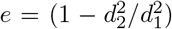, is the eccentricity of the SPB. Noting that the diameter of a MT is 2*d_MT_* = 25 nm [6], the maximum number of MTs that can be accommodated on each SPB is

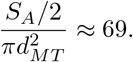

The factor of half above takes care of the fact that MTs grow from half of the area of a SPB. Finally considering both the SPBs at the two poles, we obtain a maximun number of 2 × 69 = 138 MTs which may grow within a *fission yeast* nucleus.

### Estimating typical time using extreme value statistics — derivation of Eq. (4) in the main text

Suppose, {*t*_1_,*t*_2_,…, *t*_*N*_1__} is a set of independent random variables such that each variable *t_i_* follows identical distributions, which have exponential tails at large *t_i_* with the same typical value *τ*:

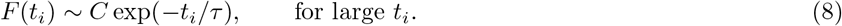

Where, *i* = 1,…, *N* and *C* is a constant. If *t_max_* represents the maximum (extreme) value from each set such that *t_max_* = *max*{*t_i_*}, then the cumulative distribution of *t_max_* becomes

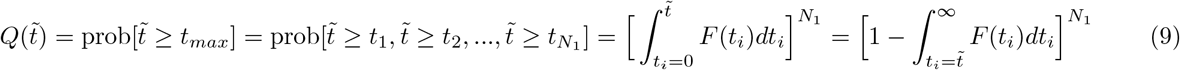

At large 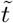, replacing *F*(*t_i_*) in Eq. (9) by Eq. (8) we get

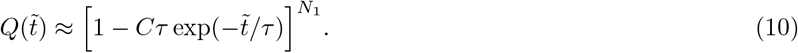

When *N*_1_ is large, 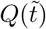 in Eq. (10) can be approximated as follows

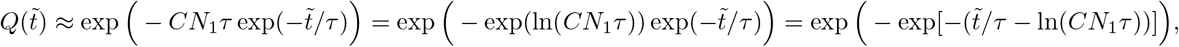

and it finally converges to the cumulative Gumbel distribution [7]

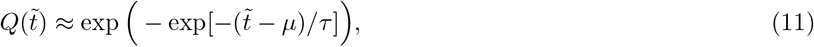

with *μ* = *τ* ln(*CN*_1_*τ*).

It is well-known that the variance *σ*^2^ of the Gumbel distribution is related to the typical value *τ* as follows [7]:

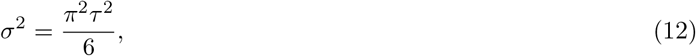

which gives typical time *τ* as

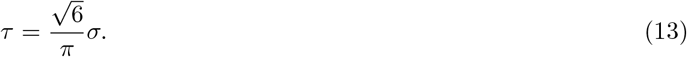

The Eq. (13) is same as Eq. (4) in the main text. In our work, we find the variance *σ*^2^ of *t_max_* from *N*_2_ sets each containing *N*_1_ sample capture times, and estimate *τ* using Eq. (13) above.

